# An *E. coli*-based platform for the production and assembly of anellovirus vectors

**DOI:** 10.1101/2024.10.30.621199

**Authors:** Rohini Prakash, Roderick Slavcev

## Abstract

Gene therapy offers immense potential for treating various diseases, including cancer, immunodeficiencies, and cardiovascular conditions. The efficacy of gene therapy (GT) largely depends on the vector used for gene delivery. Viral vectors, while effective, pose risks including insertional mutagenesis, immune responses, and high manufacturing costs. Non-viral vectors, although safer and easier to produce, often exhibit lower transfection efficiency and weaker transgene expression. This highlights the need for novel, more efficient vectors. Among emerging strategies, bacteriophages are gaining attention as promising GT delivery vehicles due to their adaptability and safety profile. Filamentous phages like M13 have demonstrated potential as targeted gene delivery vectors. This study proposes constructing a single-stranded DNA (ssDNA) phage-based vector incorporating a eukaryotic gene cassette. By leveraging Ff phage replication mechanisms in *E. coli*, the study explores encapsidating ssDNA within anellovirus capsids. These small, ssDNA viruses, known for their ability to transfect diverse tissues, offer a safer alternative to conventional viral vectors. Through successful expression and assembly of anellovirus capsid proteins, ssDNA viral particles were produced *ex vivo*. This innovative *E. coli*-based anellovirus-phagemid system provides a promising, cost-effective platform for developing next-generation viral vectors in gene therapy.

## Introduction

Gene therapy (GT) is an advanced approach in the field of medicine, aimed at treating both acquired and inherited diseases by altering gene expression. This innovative approach focuses on delivering specific genetic material into a patient’s cells to correct or modify the expression of targeted genes, thereby addressing the root cause of a disease rather than merely treating symptoms. The success of gene therapy hinges heavily on selecting an effective gene delivery vector—a mechanism that transports the therapeutic gene into desired cells. An ideal vector should possess several characteristics, such as high specificity to target cells, efficiency in gene delivery, minimal recognition by the immune system, scalability for production in high concentrations, non-allergenicity, and the ability to maintain transgene expression for the duration of the treatment cycle [1].

Gene therapy vectors can be classified into two main types: viral and non-viral vectors. Viral vectors, derived from viruses engineered to carry therapeutic genes, are commonly used due to their high efficiency in gene transfer and stable transgene expression. They have the ability to infect a wide range of cell types, which enhances their effectiveness [2]. However, viral vectors present challenges, including limited genetic material capacity, the risk of immune reactions, off-target effects, inflammation, and the potential for insertional mutagenesis, which can vary in severity depending on the virus used [2].

Conversely, non-viral vectors are generally safer, easier to produce, and less likely to induce an immune response. Despite these advantages, non-viral vectors often suffer from low gene delivery efficiency and poor transgene expression [2]. Although both vector types have favorable traits for gene therapy, no single vector completely fulfills the ideal requirements, which remains a significant obstacle in genetic medicine.

Consequently, researchers are exploring novel approaches to develop vectors that enhance transfection efficiency, transgene expression, safety, and scalability. One promising avenue involves the use of anelloviruses as potential gene delivery vectors.

Anelloviruses are a diverse group of viruses found abundantly in human and animal viromes, meaning they coexist with hosts without causing disease. This suggests a stable relationship with the host immune system, presenting a unique advantage as gene delivery vectors [3]. According to the International Committee on Taxonomy of Viruses (ICTV), the Anelloviridae family comprises 30 genera, among which alphatorquevirus, betatorquevirus, and gammatorquevirus are the most thoroughly studied [4,5]. Notably, torque tenovirus (TTV), part of the alphatorquevirus genera, along with torque teno mini virus (TTMV) and torque teno midi virus (TTMDV), which belong to the betatorquevirus and gammatorquevirus genera, respectively, have garnered considerable interest due to their structural and genetic similarities [6,7].

Anelloviruses are characterized by non-enveloped icosahedral structures, as evidenced by electron microscopy imaging, and contain single-stranded negative-sense circular DNA (ssDNA) genomes. These genomes, which range in size from 2.0 to 3.9 kb, replicate via rolling circle mechanisms, aided by a unique structure in their untranslated region (UTR) that forms hairpins. The anellovirus genome also contains overlapping open reading frames (ORFs) encoding proteins essential for viral function [4,6,7,8]. Among these proteins, ORF1, which exhibits DNA-binding properties due to arginine repeats in its N- terminal region, appears to play a key role in DNA packaging, akin to the Cap proteins of circoviruses [9]. Additionally, ORF2 is thought to be a regulatory protein that helps the virus evade the host immune response [10,11]. Anelloviruses are unique in that they pervade various cells and tissues, indicating that they do not have a strong preference for a specific cell type. This broad distribution, combined with their presence in exosomes, protects them from the immune system’s neutralizing efforts and enhances their infectivity. Exosomes not only shield anelloviruses from the innate immune response but also promote infectivity in non-permissive cells, potentially enabling redosability, as these viruses can suppress T-cell mediated adaptive immune responses [12,13].

Despite their potential, anelloviruses are still poorly understood, and extensive research is required to fully explore their capabilities as gene delivery vectors. Currently, no effective system or animal model exists to cultivate anelloviruses *ex vivo*, posing a barrier to their widespread adoption. Additionally, the high cost of manufacturing viral vectors remains a challenge in the field. The limited *in vitro* studies that have been conducted have shown that anelloviruses may be an ideal vector for gene therapy. We propose that minimal circular ssDNA molecules encoding a transgene cistron of interest and carrying a packaging signal for anellovirus can be produced and purified as M13 miniphagemids from host *E. coli* cells. Based on virus distribution, stability, and DNA packaging efficiency, TTV19 was chosen over other known species of anellovirus for capsid protein production and the use of its region which contains the DNA binding signal. Anellovirus capsid proteins with His-tags can be produced in *E. coli*. These capsids, combined with M13-processed single-stranded circular DNA vectors can self-assemble *in vitro*. This method is a novel approach to produce anelloviral vectors.

## Materials and Methods

### Strains and Vectors

*E. coli* BL21 strains were used in the production of anellovirus capsid proteins and *E. coli* JM109 was used for DNA vector plasmid amplification and production of M13 phagemids that package the ssDNA containing anellovirus packaging signal. All the plasmids used in the study are listed in Supplemental Table 1. Bacterial strains were cultured in Luria–Bertani (LB) liquid medium, supplemented with the relevant antibiotic as required.

### Transformation of *E. coli* cells with plasmids

Calcium-competent *E. coli* cells were transformed by vectors and plated on LB plates + selective antibiotic overnight and plates were observed for growth the next day. The transformed colonies were maintained on LB plates + antibiotics. Subsequently, a plasmid extraction was carried out to confirm the transformation of the cells using the Monarch Plasmid Miniprep Kit (New England Biolabs) and digested with BamHI HF (New England Biolabs).

### Protein production and extraction

The transformed *E. coli* cells were grown in LB + Kn broth until the culture reached an optical density of A_600_ = 0.6. Once the culture reached an optical density of A_600_ = 0.6, the cells were induced with 1mM IPTG and were further grown for 4-5 h for protein production. The cells were pelleted by centrifuging at 10,000 x g for 10 min. The pellet was washed with cold PBS twice. The collected cell pellet was extracted using a cell extraction buffer from Invitrogen. 1 ml of extraction buffer was added per 10^8^ cells. The cells were lysed in a cell extraction buffer for 30 min on ice, with vortexing at 10 min intervals. The lysed cells were centrifuged at 10,000 x g for 10 min. at 4° C. The supernatant was collected for further analysis.

### SDS-PAGE and Western Blot

The protein detection component of this study was completed by conducting SDS- PAGE followed by Western blotting. Approximately 10 ug of protein was mixed with 10 µL of 4X laemmeli buffer and heater at 95° C for 5 min. using a dry block heater. 20 µL of samples and 3.5 µL of BLUeye prestained protein ladder (Frogga Bio Scientific Solution) were loaded into the wells. The gels were run at 90 V for the first 15 min and then at 120 V for approximately 1 h in 1X running buffer.

Protein from the gel was transferred onto 0.45 µm nitrocellulose membrane (Bio- Rad) using a Trans-Blot SD Semi-Dry Transfer cell (Bio-Rad) at a voltage of 25 V for 30 min. The membrane was then blocked using 5% skimmed milk in Tris-HCl buffered saline/ Tween-20 (TBS-T) for a minimum of 1 h while shaking. After blocking, it was washed thrice with TBS-T and incubated overnight with a primary 6X-His Tag monoclonal antibody (3D5) HRP (Thermo Fisher Scientific) at a concentration of 1 µg/ml. The membrane was then rewashed 3X with TBS-T and then treated with secondary antibody Goat anti-Mouse IgG (H+L) HRP from (Invitrogen) diluted with blocking buffer for 1 h while shaking. Membrane was then washed again, three times with TBS-T. Protein presence was visualized by adding Pierce^TM^ 1-Step^TM^ Ultra TMB blotting solution (Thermo Fisher Scientific) to the membrane. The membrane was then covered in foil and incubated for 30 min at RT while shaking.

### Protein Purification

The over-expressed protein of interest was purified using His60 Ni superflow resin and gravity column kit (Clontech Laboratories Inc.). The His60 Ni Gravity column (1 mL resin) and all the buffers were equilibrated to the working temperature of 4° C. The column was washed with 5-10 column volumes of His60 Ni equilibration buffer. The extracted protein sample was added to the column and was allowed to bind the resin by slowly inverting the column and incubating the column at 4° C for 1 h. After incubation, the column was installed in a vertical position, and the resin was allowed to settle. Once the resin was settled, 1 mL fractions were collected. The column was then washed with 10 column volumes of His60 Ni equilibration buffer followed by 10 columns volumes of His60 Ni wash buffer. The targeted protein was then eluted with approximately 10 column volumes of elution buffer and 1 mL fractions were collected.

### Protein quantification

The Pierce BCA Protein Assay Kit (Thermo Fischer Scientific) was used to determine the amount of protein present in the purified fractions. The albumin standards were prepared at concentrations ranging from 0 to 2000 µg/ml. Next, 10 µL of the standards and samples were plated on a non-treated Nunc 96 flat-well plate (Thermo Fischer Scientific). 200 µL of working reagent (Reagent A and Reagent B at 50:1 ratio) was then added to each well, mixed with the samples, and incubated at 37° C for 30 min. The absorbances were subsequently read at 562 nm using the plate reader (Biotek). The protein concentration within the purified samples were calculated using the line of best-fit equation determined from the albumin standards.

### Phagemid production

A fresh colony of *E. coli* JM109 containing plasmids M13SW8 and TTV19IR_CMV_gfp_M13 was grown in LB + Kn broth overnight. The next day, the culture was centrifuged (8,000 X g, 10 min) to separate the bacterial pellet from the supernatant containing the phage lysate.

### Purification of phagemid

The M13 phagemid lysate was then filtered through a 60 mL syringe using a 0.22 µm filter into a fresh 50 ml centrifuge tube. The lysate was subjected to polyethylene glycol (PEG) precipitation. The filtered lysate was concentrated by adding 1/5^th^ the volume in PEG/NaCl and mixed by inversion. The mixture was incubated at 4° C for 2h and centrifuged at 12,000 x g for 15 min at 4° C. The white pellet was re- suspended in cold TN buffer and a second PEG precipitation was carried out to further concentrate the phagemid. The PEG-precipitated phagemid was then treated with DNaseI (Promega) for 1 h at 37° C to remove any contaminating bacterial DNA. The concentrated phagemid was stored at 4° C.

### Phenol-chloroform extraction

The PEG-concentrated phagemid lysate was mixed with phenol (1:1, v/v) and subsequently vortexed. The mixture was centrifuged at 4° C for 5 min, and the top aqueous layer was extracted. This layer was extracted again with an equivalent volume of phenol: chloroform twice and once with chloroform. The phagemid ssDNA was precipitated overnight in 95% ethanol at -80° C. The precipitated ssDNA was washed twice with 70% ethanol, dried, and re-suspended in DNase and RNase- free water. The ssDNA was quantified using Nanodrop analysis. The ssDNA was then subjected to gel electrophoresis to confirm the size of the ssDNA.

### *In vitro* assembly of viruses

To a final concentration of 7.5 mg/mL, the capsid proteins were suspended in 5X assembly buffer (pH 7.2, 250 mM Tris-HCl buffer containing 250 mM NaCl, 50 mM KCl, and 25 mM MgCl_2_). A 1X assembly buffer solution containing 50 mM Tris- HCl, 50 mM NaCl, 10 mM KCl, and 5 mM MgCl_2_ with a pH of 7.2 was created by combining capsid protein and ssDNA in a 6:1 mass ratio (w/w) of capsid protein to ssDNA and a 4:1 (v/v) ratio of Milli-Q water to buffer. This was then incubated for ssDNA at 4°C for at least overnight before analysis and purification. The capsid proteins were suspended in an assembly buffer without ssDNA that was used as the negative control for qPCR and functional assembly. Similarly, ssDNA in the absence of protein served as an unencapsidated control for DNase digestion assays.

### DNase enzymatic digestion assay

Anellovirus capsid-assembled ssDNA and protein and/or ssDNA-only control was subjected to DNase treatment. 1 µL of DNase enzyme was added to the samples. The samples were then incubated at 37° C for 1 h. After the treatment of samples with DNase, the samples were treated with 1 µL of DNase stop solution and incubated at 65° C for 15 min. To visualize ssDNA digestion AGE was performed. The samples were stored at 4° C for further analysis.

### Iodixanol ultracentrifugation

The crude protein extract and assembled ssDNA with the crude capsid protein were subjected to iodixanol ultracentrifugation. The iodixanol gradient (10-60%) was prepared using OptiPrep^TM^ and 1X NTC solution. To prepare a step gradient, a 1.5mL layer of each concentration starting from the highest to the lowest (60% → 10%) was added to thin wall tubes for the SW41 rotor. The gradient tube was then stored at 4° C. Alternately, the samples were prepared by pelleting through ultracentrifugation in an SW41 T1 rotor at 4° C for 90 min at 26,000 rpm. The pellet was re-suspended in chilled 1X NTC solution. The samples were layered onto the iodixanol density gradient, followed by ultracentrifugation in an SW41 T1 rotor at 4°C for 150 min at 35,000 rpm. After ultracentrifugation, 12 fractions (with a volume of ∼ 1 mL each) were collected.

### Transmission Electron Microscopy (TEM)

The assembled capsid protein and ssDNA and capsid proteins were visualized using TEM. 20 µL of sample was added onto copper grids covered with carbon made by Electron Microscopy Sciences (ThermoFisher Scientific) and incubated for 1-2 min.

The sample solutions were removed from the grid using the Whatman Cellulose Filter Paper. The grid was washed by adding 40 µL of MilliQ water. The MilliQ water was then removed from the grid using filter paper. The grid was stained with 20 µL of 2% phosphotungstic acid for 1 min. The stain was removed using filter paper, and the grid was washed 5X with MilliQ water. The grid was then air-dried overnight and subsequently imaged using the Phillips CM10 transmission electron microscope at a magnification of 64 000x.

### Polymerase chain reaction (PCR)

PCR was carried out for extracted ssDNA from capsid proteins using primer sequences indicated in Supplementary Table 2. The C1000 Thermal Cycler was used for PCR amplification (Bio-Rad, Hercules, USA). The PCR reaction contained 10 µL of Phusion Flash High Fidelity PCR Master Mix (Thermo Fisher Scientific), 5 µL of sample DNA to be amplified, 0.5 µM of forward and reverse primers, and sterile water within a final reaction volume of 20 µL. Supplementary Table 3 outlines the settings applied for the PCR reaction. The PCR amplified products were then run on a 2% gel for approximately 1.5 h at 88 V to verify for amplification of the ∼ 100 bp *gfp* gene region.

### Quantification of virion particles

Virus particles were quantified using quantitative PCR (qPCR). A calibration curve for quantifying virion particles was constructed using the pTTV19IR_CMV_gfp_M13 plasmid that encodes the *gfp* gene. The *gfp* gene region was used for the amplification of the DNA samples, where 100-fold serial dilutions (gc/µl) of the plasmid were made (10^0^ -10^8^) to generate the calibration curve. For each qPCR amplification, the PCR reaction was prepared using 5 µL of PowerUp SYBR Green Mix (Thermo Fisher Scientific), 1 µL each of 500nM primer (forward and reverse), 2 µL of the DNA sample, and 1 µL of dH_2_O for a final reaction volume of 10 µL. PCR cycling conditions were as follows: 50° C for 2 min, 95° C for 2 min, followed by 40 cycles at 95° C for 15 s and 60° C for 1 min. Next, the melt curve was set for 1 cycle at 95° C for 15 s, 60° C for 1 min, and 95° C for 15 s. PCR reactions were run in triplicate on the StepOne Plus Real-Time PCR system (Applied Biosystems). The quantification cycle or threshold cycle number (C_t_) for each reaction was recorded and used in subsequent analysis. The following equation converts the mass of dsDNA to the quantity of the genome copies:

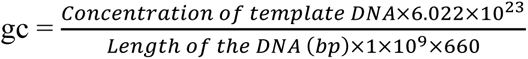

where gc is the concentration of virion genome copies (genome copies/µL).

Virus concentrations were estimated from their respective calibration curve. First, the C_t_ values from the control were plotted against the log of the known concentrations for each concentration in the dilution series. Linear regression produces a familiar equation: C_t_ = mx + b, where C_t_ is the measured threshold cycle number, m is the slope, x is the base-10, log of the concentration in gc/µL and b is the x-intercept.

PCR amplification efficiency is given by E = 10^−1/m^ – 1. The virion concentration, V can be estimated simply by the equation: V = 10^(Ct−b)/m^ × 2, where multiplication by 2 adjusts for the estimation of single-stranded products (gc/µL) from dsDNA standards. Only C_t_ measurements within the bounds of the calibration curve were used to calculate virion concentration.

## Results

### Production of TTV19 capsid protein for *in vitro* virus assembly

The possibility of TTV19 capsid protein production in a bacterial system was investigated by transforming *E. coli* BL21 with the constructed pET-28_TTV19Cap vector that expresses TTV19 ORF1 in combination with an N-terminal 6X His-tag. The production of TTV19 capsid protein in *E. coli* BL21 was evaluated using SDS-PAGE and Western blot analysis (Figure 1). TTV19 capsid protein expression was confirmed by the presence of a band at ∼75 kDa, which was the expected size. However, along with the expected capsid protein size, a band of lower molecular weight (∼50 kDa) was observed. While the reason behind this band is unconfirmed, it is likely due to proteolytic cleavage and has been observed by other investigators [14].

**FIGURE 1.**
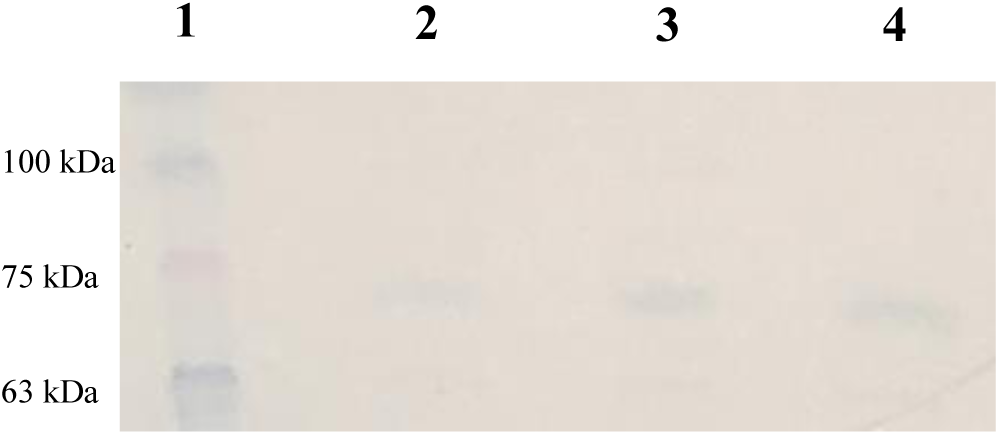
Production of TTV19 capsid protein. TTV19 capsid protein was produced via *E. coli* BL21 cells. The expressed proteins in the cell lysate were detected using western blot. A) Lane 1 is protein molecular weight ladder in kDa. B) Lanes 2, 3, and 4 shows protein bands at ∼75kDa and ∼50kDa.

### Optimization of TTV19 Capsid protein production

Once confirming that *E. coli* BL21 cells were able to produce the TTV19 capsid protein, experiments were conducted to obtain maximal capsid production within this system. The TTV19 capsid protein sequence is under the control of the *lac* operator, regulating the T7 promoter, and thus requires IPTG induction to result in high-level expression. Therefore, TTV19 capsid protein expression was evaluated following IPTG induction at various optical densities. The expression of the protein was confirmed using SDS-PAGE and Western blot analysis (Figure 2). From the analysis, it was observed that capsid proteins were expressed at all the tested optical densities by the presence of a band at ∼75 kDa. However, the maximal expression was observed at optical density A_600_ = 0.6. As such, for this study, TTV19 capsid proteins were expressed after inducing the cells with IPTG at A_600_ = 0.6. Along with proteins expressed at ∼75 kDa, protein bands of lower molecular weight were observed at ∼25 kDa. These bands were smaller than those observed in Figure 1. From this, it can be inferred that the proteins may be degraded sequentially to produce a mature virus capsid head, or that there is a proteolytic cleavage occurring in the bacterial system.

**FIGURE 2:**
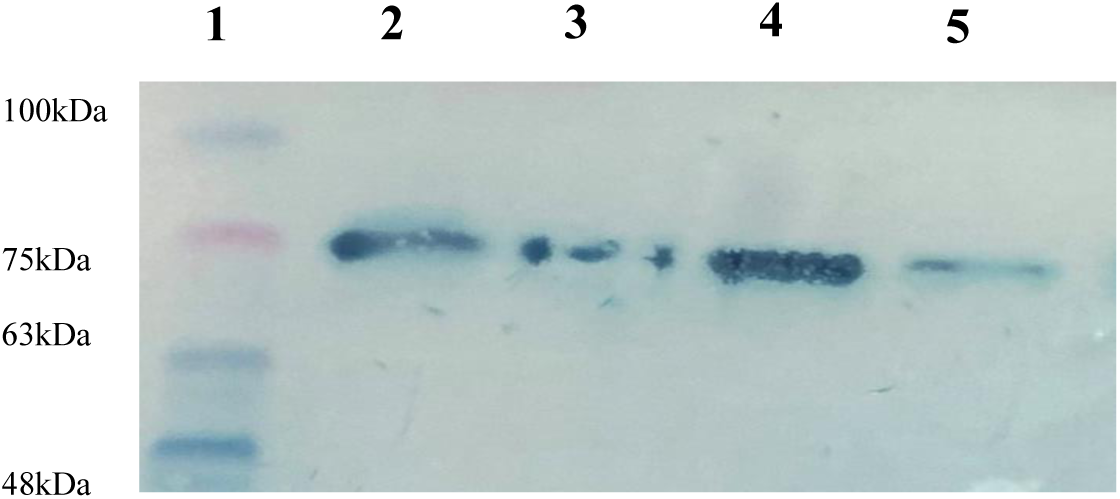
Production of TTV19 Capsid protein at different O.D. The expressed proteins in the cell lysate were detected using western blot. A) Lane 1 is protein molecular weight ladder in kDa. B) Lane 2 is IPTG induction at A_600 =_ 0 which showed the presence of protein band at 75kDa. C) Lane 3 is IPTG induction at A_600 =_ 0.4 which showed the presence of protein band at 75kDa. D) Lane 4 is IPTG induction at A_600 =_ 0.6 which showed the presence of two protein bands at 75kDa and 25kDa. E) Lane 5 is IPTG induction at A_600 =_ 0.8 which showed the presence of two protein bands at 75kDa and 25kDa.

### Purification of expressed TTV19 capsid protein

Once the TTV19 capsid proteins were optimally expressed in *E. coli* BL21, the expressed proteins were purified for *in vitro* assembly of virus particles using the Ni- NTA purification column, as the capsid protein sequence has a 6X His-tag at the N- terminal end of the capsid protein. After purification, the eluted protein samples were subjected to SDS-PAGE and Western blot analysis (anti-His Ab) to detect the presence of purified capsid proteins, as shown in Figure 3. From Figure 3, it was observed that the proteins were successfully purified, although, somewhat unexpectedly, the molecular weight of the purified protein was ∼25 kDa, which is smaller compared to the expected protein molecular weight (∼75 kDa). From Figure 3A, it was observed that the expressed proteins in crude extracts produced lighter protein bands at ∼75 kDa, along with another protein band at ∼50 kDa. However, once the proteins were purified, proteins that were ∼25 kDa in size were observed in elution fractions 1, 2, and 3 following purification. While the origin of the smaller His^+^ protein is unknown, it is quite likely that the mature 75 kDa ORF1 protein has been further processed to generate a 25 kDa product. Whether this processing is specific to capsid maturation and resulted in a mature virus capsid head or if proteolytic cleavage occurred in the bacterial system is not yet known. Therefore, the purified protein and the crude extract containing the capsid proteins were both used for further *in vitro* assembly of virus particles.

**FIGURE 3:**
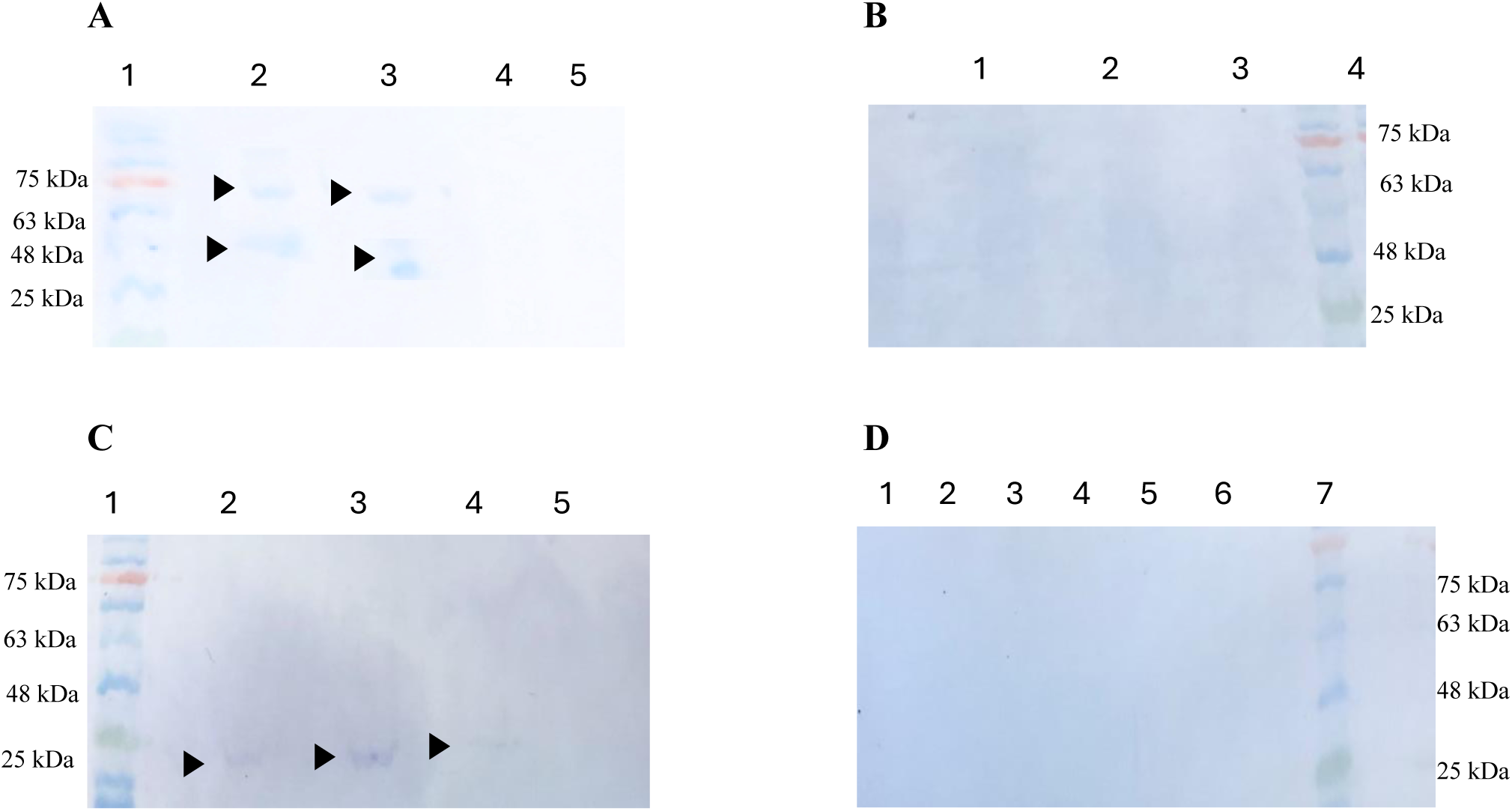
Purification of His-Tagged TTV19 capsid protein. A) Lane 1 is protein molecular weight ladder, Lane 2& 3 is crude extract containing TTV19 capsid protein, Lane 4&5 is BW1 and BW2. B) Lane 1,2, & 3 are BW5, BW4, and BW3 respectively, Lane 4 is protein molecular weight ladder. C) Lane 1 is protein molecular weight ladder, Lane 2,3,4,&5 is protein samples after elution, E1, E2, E3, and E4 respectively, D) Lane 1,2,3,4,5,&6 are eluted proteins samples E10, E9, E8, E7, E6, and E5 respectively, and Lane 7 is protein molecular weight ladder. Key: CE = Crude protein extract containing TTV19 capsid protein, BW = Samples collected by washing the columns before eluting the sample, and E = Sample elution from the column.

Next, the concentration of purified protein samples was determined for *in vitro* virus assembly. A standard curve as shown in Supplementary Figure 1 was obtained for albumin concentrations. From the standard graph, a linear relationship of y = 0.000x + 0.0211 was observed and R^2^ value of 0.9986. The unknown concentration of purified TTV19 capsid proteins was determined using this formula. The average concentration of the protein sample was found to be 39.4985 ± 9.148 µg/mL. The purified TTV19 capsid protein samples were used for *in vitro* assembly of ssDNA.

### Production of ssDNA containing the TTV19 DNA binding signal using M13 replication system

The second component of the viral construct focused on the production of ssDNA for *in vitro* assembly of virus particles. This was achieved by transforming *E. coli* JM109 cells by pTTV19 IR_CMV_gfp_M13 (for packaging of ssDNA) and the helper plasmid (pM13SW8) for M13 phagemid production.

### Production, purification, and quantification of M13 phagemid containing the TTV19 DNA binding signal

To obtain the phagemid titer, qPCR was performed where a segment of the *gfp* gene region was amplified. The average phage concentration was found to be 1.53 x 10^9^ ± 1.66x 10^8^ phage/mL.

### Extraction of ssDNA from M13 phagemid for *in vitro* assembly

After M13 phagemids were produced and purified, ssDNA was extracted from M13 phagemids. The ssDNA is ∼1.6 kb in size but is expected to appear on the gel as half the size ∼0.8 kb since the DNA sample is single-stranded and will run faster compared to the dsDNA ladder. The extracted ssDNA was visualized using AGE and appeared on the gel at a size of ∼ 0.8kb as predicted (Figure 4). This extracted ssDNA will be used for *in vitro* assembly.

**FIGURE 4:**
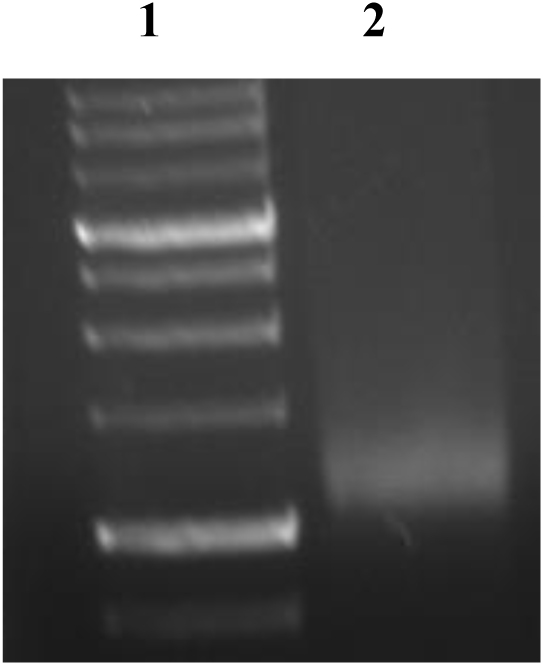
ssDNA obtained from M13 phagemid carrying the gene cassette. A) Lane 2 is DNA ladder. B) Lane 1 shows ssDNA obtained from M13 phagemid which is ∼1.6kb. Since the DNA is ss it will run faster, and the DNA should be ∼0.8kb as observed above.

### *In vitro* assembly of ssDNA containing the gene of interest and TTV19 intergenic region, and TTV19 capsid protein

The next step of the study involved the *in vitro* assembly of capsid proteins and the extracted ssDNA. This was achieved by assembling His-tag purified TTV19 capsid protein and ssDNA. Along with His-tag purified capsid proteins, a crude extract containing TTV19 capsid protein and ssDNA were also assembled *in vitro*. The assembly of virus particles and packaging of ssDNA was confirmed using the experiments described below.

### Iodixanol ultracentrifugation of crude extract containing TTV capsid protein and assembled crude extract containing TTV19 capsid protein and ssDNA

Once the crude extract containing TTV19 capsid proteins assembled ssDNA, the assembled products were subjected to iodixanol ultracentrifugation for purification of the assembled virus particles. Along with the assembled capsid protein sample, a crude extract containing only TTV19 capsid was also subjected to iodixanol ultracentrifugation. The purified samples were then subjected to SDS-PAGE and Western blot to detect the presence of purified capsid protein in the fraction. The purified proteins for both samples were obtained in fraction 10 at a band size of ∼25 kDa, as shown in Figure 5 and Figure 6. For further study, fraction 10 of the iodixanol centrifuged samples of crude extract containing TTV19 capsid protein and fraction 10 of the iodixanol centrifuged samples of assembled ssDNA with crude extraction containing TTV19 capsid protein, were used.

**FIGURE 5:**
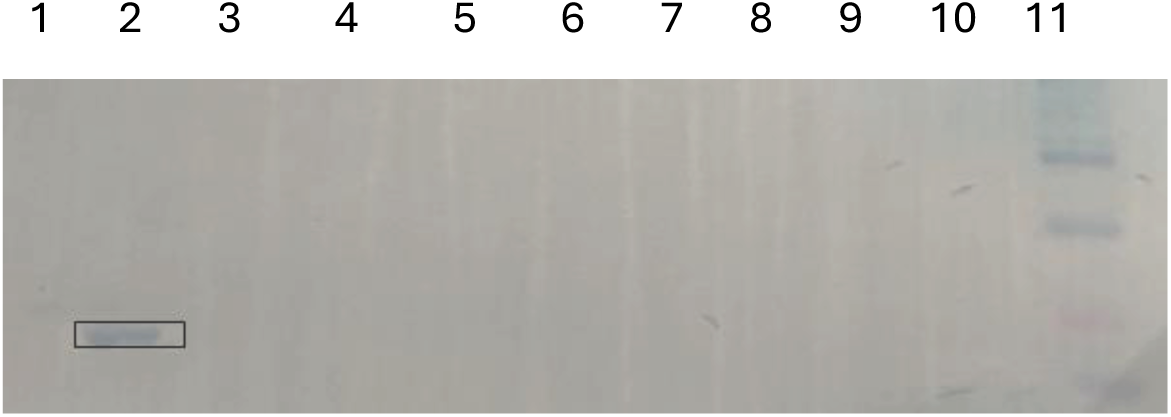
Iodixanol ultracentrifugation of crude extract containing TTV19 capsid protein. Lane 1 – 10 is F11 to F1 respectively, and Lane 12 is protein molecular weight ladder Key: F = Fraction.

**FIGURE 6:**
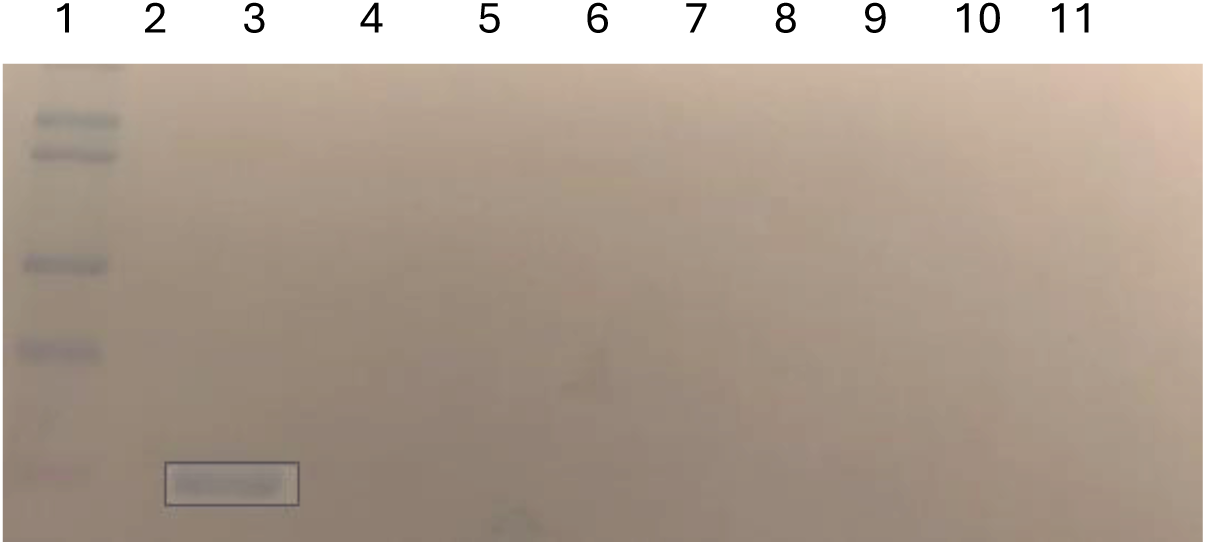
Iodixanol ultracentrifugation of in vitro assembled ssDNA and crude extract containing TTV19 capsid protein. Lane 1 is protein molecular weight ladder, and Lanes 2 – 12 is F11 to F1 respectively. Key: F = Fraction.

### Transmission electron microscopy (TEM) for *in vitro* assembled TTV19 capsid protein and ssDNA containing the gene of interest

After assembling capsid proteins and ssDNA *in vitro,* the samples were examined by TEM to try to identify any present assembled virus particles. TTV19 virus is ∼50 nm and assembles as an icosahedral head. TEM images for crude extract containing TTV19 capsid protein showed the presence of what seemed to be icosahedral heads approximately ∼50 nm in size (Figure 7 A & B). While not particularly conclusive, this finding supports the hypothesis that the capsid proteins are assembling into virus- like particles (VLP) in the bacterial system.

**FIGURE 7:**
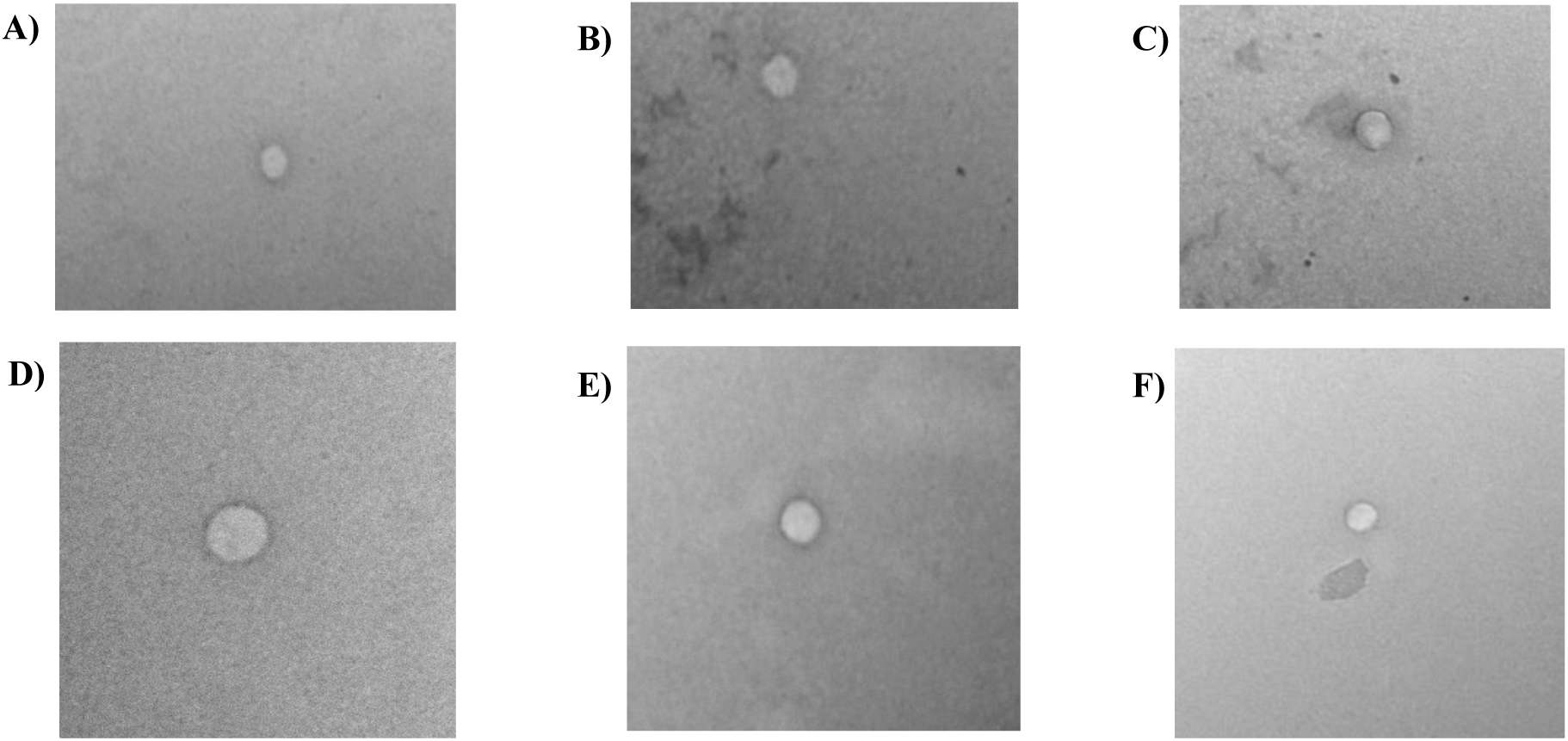
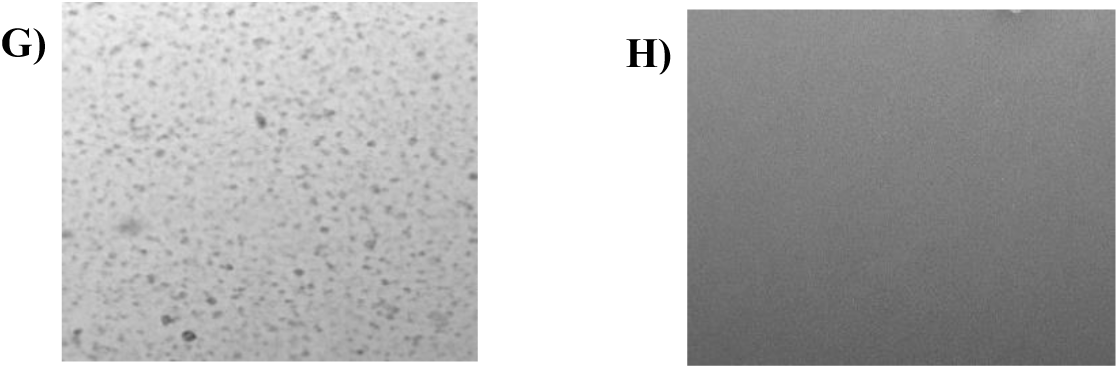
TEM images for crude extract containing TTV19 extract samples. A&B: Crude extract containing TTV19 capsid proteins; C&D: assembled crude extract containing TTV19 capsid protein and ssDNA; E&F: DNase treated assembled crude extract containing TTV19 capsid protein and ssDNA; G: Proteinase K treated assembled crude extract containing TTV19 capsid protein and H: ssDNA treated with DNase enzyme.

Both assembled crude extract containing TTV19 capsid protein and ssDNA, as well as assembled crude extract containing assembled capsid protein and ssDNA treated with DNase, indicated the presence of icosahedral heads of the correct size (Figure 7 C, D, E, & F), however, they were few in number. The negative controls for both extracts using Proteinase K show the presence of an icosahedral heads which demonstrates that the virus particles are assembling (Figure 7 G & F). To confirm the assembly of virus particles, further testing is required using more conclusive methods. Along with crude extracts, His-tag purified TTV19 capsid proteins were also assembled *in vitro* and were subjected to TEM. When the His-tag purified TTV19 capsid proteins were analyzed using TEM, no VLP was observed which differs from the observed results for crude extract containing TTV19 capsid proteins (Figure 8A). This indicated that the His-tag purified proteins don’t assemble on their own; this could be observed due to low concentrations of viral capsid proteins. Next, we subjected assembled His-tag purified capsid protein and ssDNA (Figure 8B). It was found that very few assembled into viral particles. The assembled His-tag purified TTV19 capsid protein treated with DNase was also subjected to TEM analysis, which showed the presence of icosahedral heads of the proper size (Figure 8C & D). The absence of an icosahedral head for Proteinase K-treated samples and ssDNA digested with DNase enzyme (Figure 8E & F) validates that the viral particles might be assembling to form icosahedral particles. The results show that the viruses might be assembling into icosahedral heads but whether the TTV19 capsid proteins are packaging ssDNA could not be evaluated by this method alone, and as such, further testing was required to confirm DNA encapsulation in TTV19 capsids.

**FIGURE 8:**
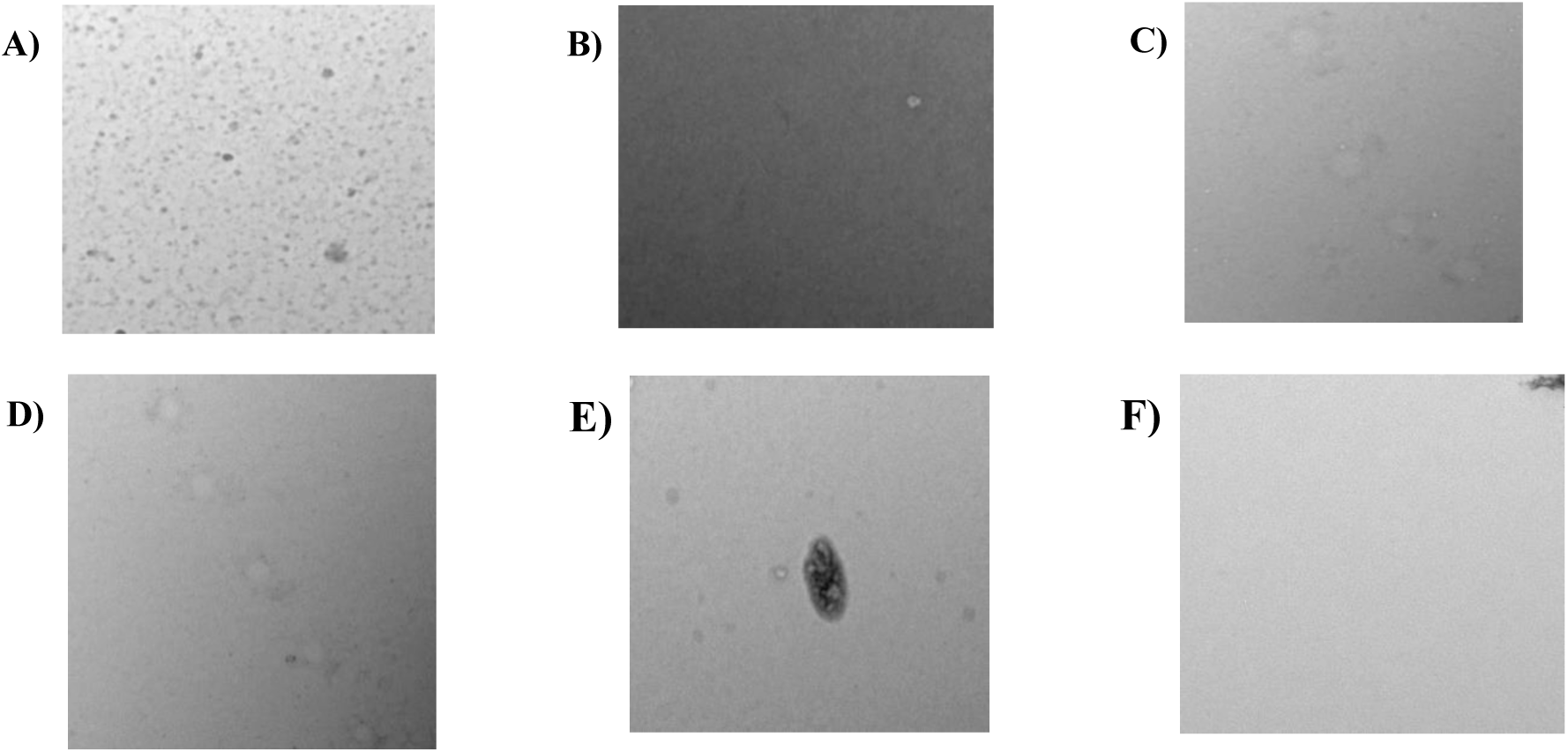
TEM images His-Tag purified TTV19 extract samples. A: His-Tag purifiedTTV19 capsid proteins; B: assembled His-Tag Purified TTV19 capsid protein and ssDNA; C&D: DNase treated assembled His-Tag purified TTV19 capsid protein and ssDNA. E: Proteinase K treated assembled His-Tag Purified TTV19 capsid protein and ssDNA; and F: ssDNA treated with DNase.

### Detection of packaging of ssDNA by TTV19 capsid protein *in vitro* by PCR

Next, to confirm that the capsid proteins are assembling ssDNA *in vitro,* PCR was performed to amplify a segment *gfp* gene region of ∼100 bp carried by the DNA vector. Following PCR amplification, the amplified DNA fragment (∼100 bp) was visualized by AGE (Figure 9). The ssDNA extracted from an assembled crude extract containing TTV19 capsid protein and ssDNA exhibited *gfp* gene amplification (∼100 bp). This indicates that the capsid proteins present in crude extract in assembling ssDNA *in vitro* (Figure 9A). On the other hand, ssDNA extracted from assembled His-tag purified capsid protein, and ssDNA did not show any amplification of the *gfp* gene (Figure 9B) which might indicate that the capsid proteins are not packaging ssDNA but assembling to form VLPs. To further prove this, more sensitive and accurate testing such as DNA quantitative methods should be performed.

**FIGURE 9:**
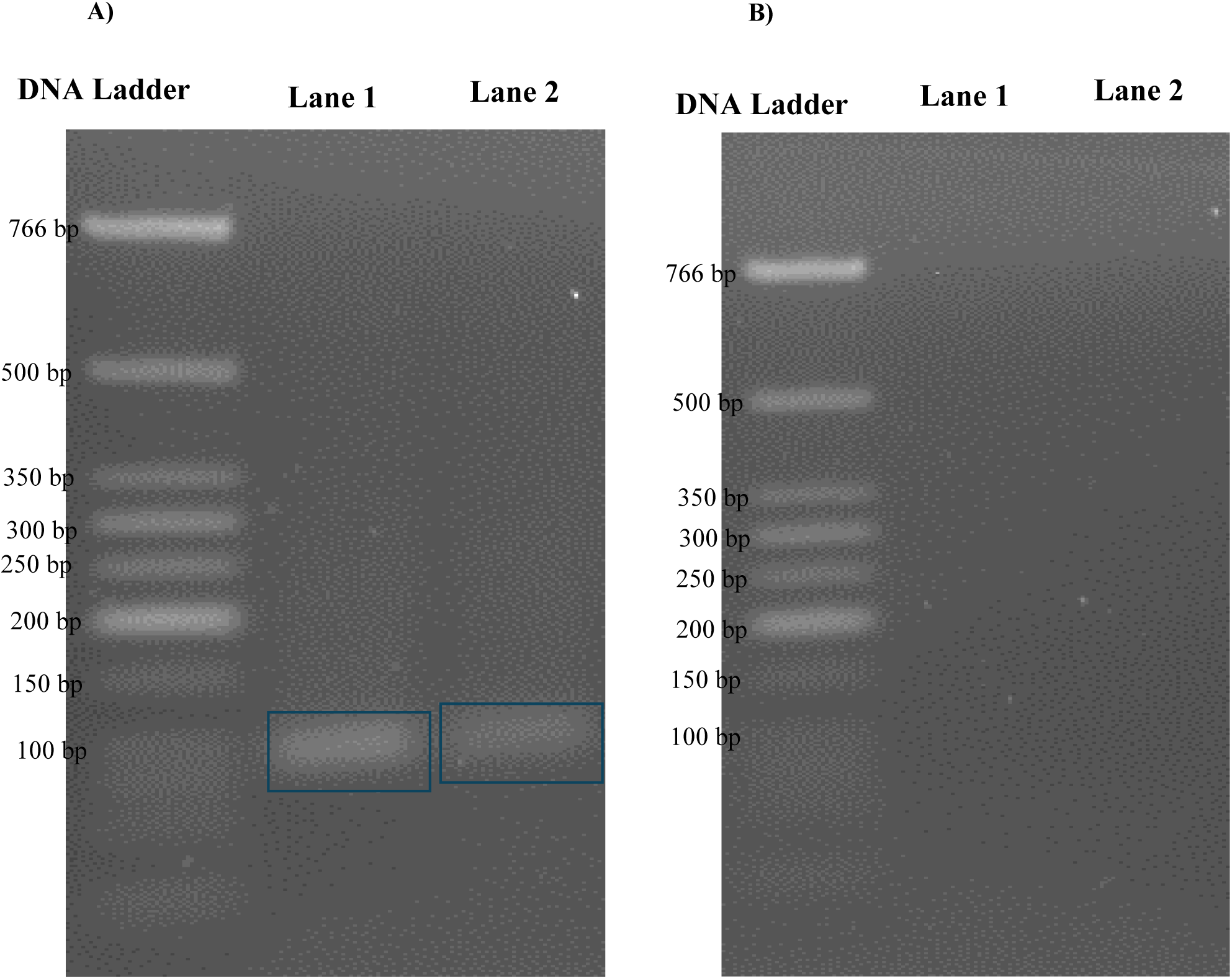
1.8% gel showing PCR amplification of gfp gene region. A: PCR amplification of the *gfp* gene from assembled crude extract containing TTV19 capsid protein and ssDNA. B: PCR amplification of the *gfp* gene from assembled His-Tag purified TTV19 capsid protein and ssDNA.

### Confirmation of packaging of ssDNA by TTV19 capsid protein and determination of concentration of TTV19 virion particles

Capsid proteins from crude extract were able to assemble ssDNA *in vitro,* while His- tag purified capsid proteins were not able to assemble ssDNA *in vitro.* To further validate the results, the DNA quantification approach of quantitative PCR (qPCR) was carried out to confirm ssDNA packaging by capsid proteins *in vitro.* This approach can also be used to determine the concentration of assembled virus particles *in vitro*. To estimate virus concentration, the standard calibration curve was generated using the plasmid pTTV19 IR_CMV_gfp_M13 (Supplementary Figure 2). Once qPCR was run on the samples, it was observed based on the amplification curves as well as the threshold cycle number (Table 1), that the ssDNA that was generated by the M13 processing was successfully packaged by crude extract containing TTV19 capsid proteins as well as crude extract containing His-tag purified TTV19 capsid proteins. ssDNA was extracted from the assembled capsid protein from the crude extract using heat treatment and phenol-chloroform, it was then successfully amplified, indicating that the DNA was likely packaged by the crude extract containing capsid protein. As expected, ssDNA treated with DNase did not show any amplification. Using the threshold cycle number, the concentration of virus particles was determined as shown in Table 1. Next, we extracted ssDNA again, using heat and phenol-chloroform, from the assembled His-tag-purified capsid protein. The DNA produced was successfully amplified, indicating that the ssDNA was likely packaged by capsid protein within the crude extract. The extracted ssDNA treated with DNase did not show any amplification in this case, as expected. This result was similar to what was obtained for the crude extract containing capsid proteins and the virus concentration was similarly determined (Table 1).

**TABLE 1:** C_t_ Values and virion concentration for the qPCR amplified samples.

| Test Sample | C<sub>t</sub> Value | Virion concentration<br>(No. of virus particles/μl) |
| --- | --- | --- |
| ssDNA + Crude extract containing TTV19 capsid protein → DNase Treatment → Heat → DNase | 30.43 ± 0.97 | NA |
| ssDNA + Crude extract containing TTV19 capsid protein → DNase Treatment → Heat | 20.80 ± 2.31 | 1.2 x 10<sup>4</sup> ± 1.82 x 10<sup>2</sup> |
| ssDNA + Crude extract containing TTV19 capsid protein → DNase Treatment → Phenol-chloroform extraction → DNase | 32.85 ± 2.90 | NA |
| ssDNA + Crude extract containing TTV19 capsid protein → DNase Treatment → Phenol-chloroform extraction | 23.68 ± 0.89 | 5.73 x 10<sup>4</sup> ± 5.47 x 10<sup>2</sup> |
| ssDNA + His-Tag Purified TTV19 capsid protein → DNase Treatment → Heat → DNase | 31.49 ± 0.47 | NA |
| ssDNA + His-Tag purified TTV19 capsid protein → DNase Treatment → Heat | 18.41 ± 0.58 | 3.01 x 10<sup>4</sup> ± 1.23 x 10<sup>3</sup> |
| ssDNA + His-Tag purified TTV19 capsid protein → DNase Treatment → Phenol-chloroform extraction → DNase | 31.93 ± 0.17 | NA |
| ssDNA + His-Tag Purified TTV19 capsid protein → DNase Treatment → Phenol-chloroform extraction | 17.94 ± 0.25 | 4.19 x 10<sup>4</sup> ± 8.48 x 10<sup>3</sup> |
| ssDNA + DNase | 31.89 ± 0.57 | NA |

## Discussion

Anelloviruses are referred to as ‘orphan viruses’ because they lack disease association, however, they are an omnipresent and commensal part of the human microbiome.

These viruses circulate throughout our tissues unimpeded by antibodies due to their associated exosomes, which also aid the virus to infect non-permissive cells. The use of bacterial systems has become more popular for virus-like particle (VLP) production as this system is i) cost-effective; ii) easy to scale up; iii) easily manipulated; iv) offers high-level expression; v) offers a faster growth rate; vi) genetically stable; and vii) confers simplicity of expression. To produce anellovirus capsid protein, TTV19 was used as a reference sequence because all the gene sequences for this virus have been identified, and it is one of the most abundant anellovirus species found in the human body [15]. The *E. coli* cell system has been successfully used to assemble a variety of VLPs. Interestingly, we did not find that the His-tag posing any issues to TTV19 capsid assembly, suggesting this may be a scalable and simplistic approach to capsid production. The TTV19 capsid expressed in *E. coli* weighs ∼75 kDa when unprocessed. While this was observed initially, the protein was then further processed to a lower molecular weight, which was determined to be ∼25 kDa upon purification. Proteolysis occurs at the C-terminus of the capsid protein and does not play an important role in VLP formation. It was suggested that proteolysis of the ORF1 C- terminus may be a natural part of the anellovirus formation. Similar results were obtained by a group of researchers while studying the structure of a virus-like particle (VLP) derived from a betatorquevirus isolate called LY1 [14]. It was found that ORF1 of TTV19 had several enzymatic cleavage sites although none of the identified enzymes would have generated the cleavage pattern. Since the TTV19 capsid protein was produced in *E. coli* BL21, a protease-deficient cell line, it is unlikely that an *E. coli* protease was responsible for the cleavage, thus the reason for the cleavage remains unknown. The final 25 kDa protein does appear to form capsids, so this may be a natural process that is occurring intrinsically within the protein. It was found that production of a synthetically originated ssDNA could be packaged by TTV19 capsid protein *in vitro*. This was achieved using the M13 phage replication system using the helper plasmid, M13SW8, and the DNA vector plasmid, pTTV19 IR_CMV_gfp_M13 and produced *in vivo* in *E. coli* host cells.

The helper plasmid encodes for the proteins necessary for M13 phage replication, transcription, and assembly of M13 phagemids intracellularly in bacterial systems [16,17]. The M13 genome consists of an origin region that has nucleotide sequences for initiation and termination of replication. While the ssDNA to be packaged by M13 phagemids was produced using a DNA vector plasmid, the M13 origin of replication was separated and flanked by a mammalian transgene expression cassette, isolating it from the bacterial backbone [16,17]. This permits the separation of the M13 initiation sequence from the termination sequence and the ability to clone a gene cassette of interest between the initiation and termination gene sequences without any intervening prokaryotic sequences or helper phage contamination [16,17]. Using this miniphagemid approach, M13 phagemids were produced that packaged ssDNA and the TTV19 DNA binding signal. This miniphagemid ssDNA production system should be similarly applicable to synthetically reconstitute a viable anellovirus and the entire original genome sequence of TTV19 using *in vivo* generated components from *E. coli*. *In vitro* assembly of ssDNA and TTV19 capsid protein was successfully carried out.

His-tag purified TTV19 capsid proteins and crude extract containing TTV19 capsid proteins were used. Both crude extract containing TTV19 capsid protein and His-tag purified capsid protein appeared to successfully assemble ssDNA. The ssDNA was observed to be assembled by His-tag purified proteins, which were 25 kDa in size, suggesting that processing capsid proteins from ∼75 kDa to 25 kDa is necessary to produce mature virus particle formation. The concentration of virus particles produced in the study was also similar for both ssDNA assembled by crude extract containing TTV19 capsid protein and His-tag purified TTV19 capsid protein. This further demonstrates that the contents of the bacterial system do not hinder virus assembly and may in fact fully support it, as well as fully supporting the production and complete synthetic assembly of anelloviruses *in vivo* in an *E. coli* system.

### Summary and Conclusion

In this study, we hypothesized that ssDNA molecules encoding a transgene cistron of interest and carrying a packaging signal for anellovirus can be produced and purified as M13 miniphagemids from host *E. coli* cells. In tandem, anelloviral capsid proteins can be expressed and functionally processed in *E. coli* to yield self-assembling homo- multimeric anellovirus capsids. M13-processed single-stranded circular DNA vector in combination with *E. coli*-produced capsids can self-assemble into circular ssDNA containing anellovirus capsid gene delivery vectors. From the results, it was observed that TTV19 capsid protein can be expressed in *E. coli*. These proteins seem to undergo additional processing and can functionally self-assemble into capsids of the expected size. In combination, ssDNA containing TTV19 DNA binding signal was produced in tandem using the M13 replication system and was obtained from M13 phagemids using phenol-chloroform extraction. TTV19 capsid proteins were able to successfully assemble ssDNA *in vitro*, thus, supporting the hypothesis of the study. The assembled TTV19 virion particle can be further tested for its transfection efficiency and transgene expression in mammalian cells to understand whether the nucleoprotein composition of this virus is suitable as a GT vector or perhaps further requires exosomal incorporation for effective transduction. Thus, the vector produced in this study is a promising candidate for the gene therapy platform. This approach opens new avenues to produce novel promising vectors for gene therapy platforms. The application can be further extended to overcome the production limitations of producing cost-effective and safer viral vectors.

## Data Availability

All relevant data are within the manuscript.

## Acknowledgements

Many Thanks to Deborah Pushparajah and Julia Lumini for their contribution in formatting and editing of this manuscript.

## Author Contributions

Conceptualization, R.P., and R.S.; Methodology, R.P., and R.S.; Formal Analysis, R.P.; Investigation, R.P.; Visualization: S.W.; Writing – Original Draft Preparation, R.P..; Writing – Review & Editing, R.P., and R.S.; Supervision, R.S.; Funding Acquisition, R.S.

