## Supplemental Figures and Tables for "An *E. coli*-based platform for the production and assembly of anellovirus vectors"

Supplementary Data

**SUPPLEMENTARY TABLE 1: Plasmids used in the study**

| Plasmid | Genotype | Source |
|---|---|---|
| pET-28_TTV19Cap | T7 promoter, lac operator, 6X His-tag, TEV cleavage site, TTV19 ORF1 | This Study |
| pTTV19 IR_CMV_gfp_M13 | M13 origin, TTV19 intergenic region, CMV promoter, *gfp* gene inserted between M13 initiation and termination sequence, $Amp^R$ | Wong, 2022 |
| M13SW8 | M13KO7, Packaging signal removed, $Kn^R$ | Wong, 2022 |

**SUPPLEMENTARY TABLE 2: Primers used in this study.**

| Primer | Sequence |
|---|---|
| **gfp-F Forward** | 5'CAAGATGAAGAGCACCAAAGG3' |
| **gfp-R Reverse** | 5'CGAAGTGGTAGAAGCCGTAG3' |

**SUPPLEMENTARY TABLE 3: PCR amplification protocol.**

| Cycle Step | Temperature | Time | Number of Cycles |
|---|---|---|---|
| **Initial Denaturation** | 98° C | 10 sec | 1 cycle |
| **Denaturation** | 98° C | 5 sec | 30 cycles |
| **Annealing** | 62° C | 5 sec | |
| **Extension** | 72° C | 2 sec | |
| **Final Extension** | 72° C | 1 min | 1 cycle |
| **Incubation** | 4° C | Incubate | N/A |

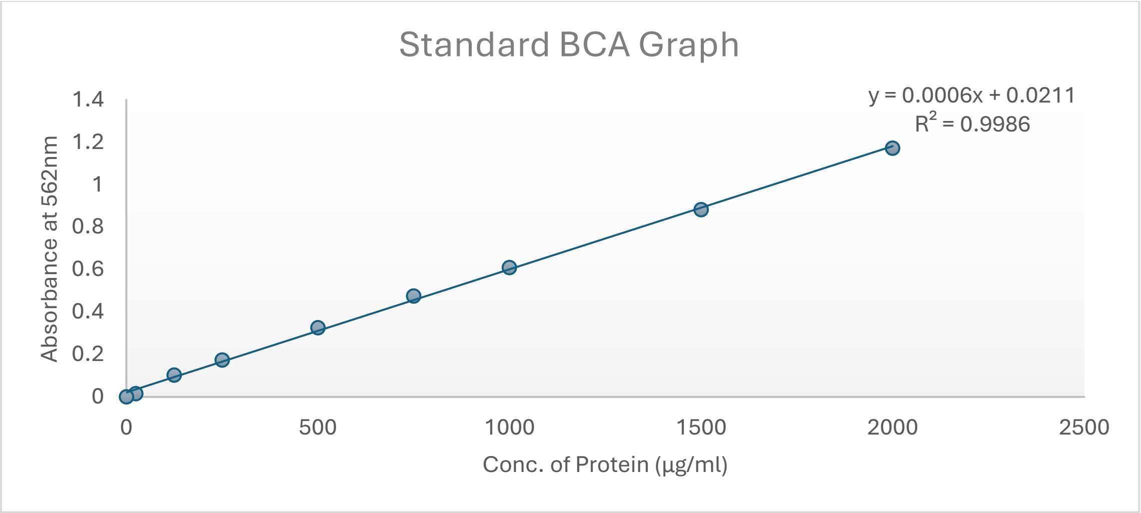


**SUPPLEMENTARY FIGURE 1: Standard BCA graph for protein concentration determination.** The standard BCA graph was obtained using albumin using concentrations ranging from 0 µg/ml to 2000 µg/ml.

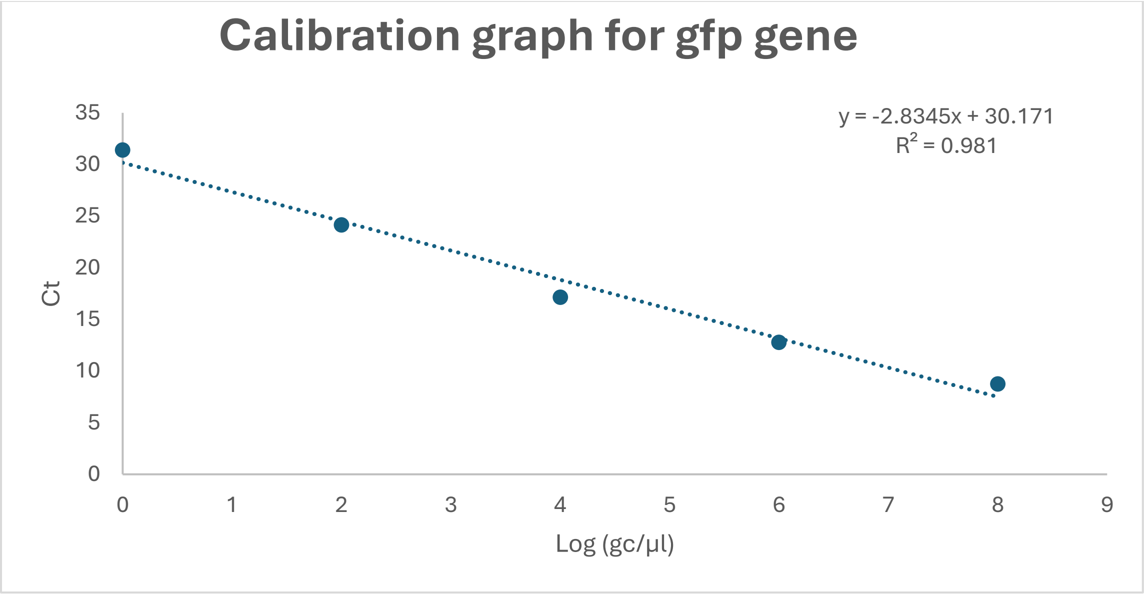


**SUPPLEMENTARY FIGURE 2: Calibration graph to estimate TTV19 virion concentration.** Representative calibration curve for *gfp* gene region measured from qPCR of pTTV19 IR_CMV_gfp_M13 plasmid.
